# Targeting PD-L1 in solid cancer with myeloid cells expressing a CAR-like immune receptor

**DOI:** 10.1101/2024.01.29.577873

**Authors:** Kayla Myers Chen, Daniel Grun, Brian Gautier, Shivaprasad Venkatesha, Michael Maddox, Ai-Hong Zhang, Peter Andersen

## Abstract

Myeloid cells are prevalent in solid cancers, but they frequently exhibit a pro-tumor phenotype, hindering cancer immunotherapy. Their abundance makes engineered myeloid cell therapy an intriguing approach to tackle challenges posed by solid cancers, such as tumor trafficking and infiltration along with tumor cell heterogenicity and immunosuppressive tumor microenvironment (TME). Solid cancers often upregulate the checkpoint molecule PD-L1 to evade immune responses. Thus, we devised an adoptive cell therapy strategy based on myeloid cells expressing a Chimeric Antigen Receptor (CAR)-like immune receptor (CARIR). The extracellular domain of CARIR is derived from the natural inhibitory receptor PD-1, while the intracellular domain(s) are derived from CD40 and/or CD3ξ. To assess the efficacy of CARIR-engineered myeloid cells, we conducted proof-of-principle experiments using co-culture and flow cytometry-based phagocytosis assays *in vitro*. Additionally, we employed a fully immune-competent syngeneic tumor mouse model to evaluate the strategy’s effectiveness *in vivo*. Co-culturing CARIR-expressing human monocytic THP-1 cells with PD-L1^+^ target cells lead to upregulation of the co-stimulatory molecule CD86 along with expression of proinflammatory cytokines TNF-1α and IL-1β. Moreover, CARIR expression significantly enhanced phagocytosis of multiple PD-L1^+^ human solid tumor cell lines *in vitro*. Similar outcomes were observed with CARIR-expressing human primary macrophages. In experiments conducted on Balb/c mice bearing aggressive 4T1 mammary tumors, infusing murine myeloid cells expressing a murine version of CARIR significantly slowed tumor growth and prolonged survival. Taken together, our results demonstrate that adoptive transfer of PD-1 CARIR-engineered myeloid cells may be an effective strategy in treating PD-L1^+^ solid tumors.

**Graphic Abstract:** 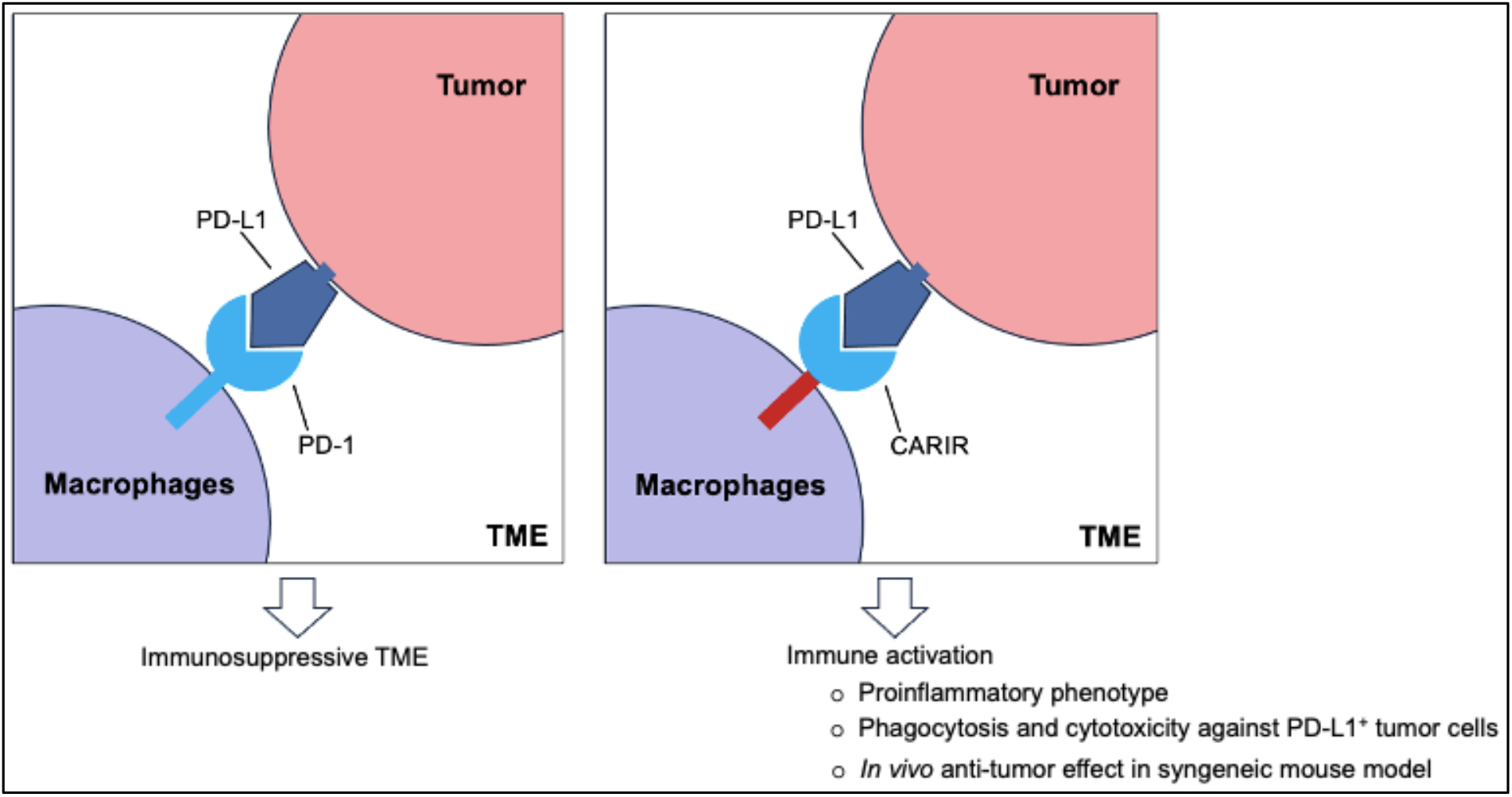

**In Brief:** We described here an adoptive cell therapy approach employing PD-L1-specific CAR-like immune receptor (CARIR) modified myeloid cells as a potential immune cell therapy strategy for treating PD-L1^+^ solid cancer.

- CARIR expression directed human THP-1 macrophages to recognize PD-L1^+^ target cells, which led to an upregulation of co-stimulatory molecule CD86 and production of proinflammatory cytokines TNF-α and IL-1β.
- CARIR expression in human THP-1 macrophages had increased % phagocytosis and killing against PD-L1^+^ tumor cells *in vitro*.
- Adoptive transfer of CARIR transduced myeloid cells in immunocompetent syngeneic mice with established aggressive 4T1 tumor significantly slowed tumor growth and prolonged survival.

## Introduction

Cancer is a leading cause of death in the US, with solid cancer accounting for nearly 90% of all cases^1^. Adoptive cell therapy using chimeric antigen receptor (CAR) T-cells has been tremendously effective against hematological malignancies such as leukemia and lymphomas, but responses in solid tumors have been elusive due to barriers for T cells infiltration, immunosuppressive tumor microenvironment (TME), and inherent tumor cell heterogeneity. Likewise, monoclonal antibody-based PD-1/PD-L1 checkpoint immunotherapies have shown durable responses only in a subset of patients^2^, indicating checkpoint inhibition alone is insufficient. Consequently, there is a high demand in developing novel and more effective therapies for solid cancer treatment.

Adoptive transfer of CAR-expressing myeloid cells, including macrophages, has emerged as a promising strategy for treating solid cancer^3,4^. In stark contrast to the limited lymphocyte infiltration, myeloid precursors are actively recruited through cancer-driven myelopoiesis. As a result, macrophages usually are the most abundant immune cells in solid cancer and can account for up to 50% of the tumor volume^5^. Being the most functionally versatile innate immune cells in the body, M1 type macrophages can destroy tumor cells themselves and orchestrating an inflammatory adaptive antitumor immune response. However, under the influence of tumor- and non-tumor derived factors in the TME, tumor associated macrophages (TAMs) are often polarized towards an anti-inflammatory, M2-like state that promote tumor growth and metastasis^6^. Notably, while the programmed cell death protein 1 (PD-1) is best known as the immune checkpoint receptor expressed on activated T cells, PD-1 expression is associated with TAMs as well^7^, indicating that PD-1/PD-L1 checkpoint therapy may exert its effect, in part, via TAMs^7,8^. Since the programmed cell death ligand 1 (PD-L1) is frequently overexpressed in solid cancer as a common pathway of immune evasion^9^, it makes PD-L1 a potential target candidate for adoptive myeloid cell therapy.

By designing a PD-L1 specific CAR-like immune receptor (CARIR) for myeloid cells, we demonstrate how myeloid cells could be redirected to targeting PD-L1^+^ solid cancer. Antitumor potency of CARIR modified human macrophages was evaluated by phagocytic activity against PD-L1^+^ tumor cells *in vitro* and using a syngeneic tumor mouse model *in vivo*. Our results demonstrate that CARIR-expressing macrophages have increased phagocytotic activity against PD-L1^+^ cancer cells *in vitro* and that systemic delivery of CARIR-expressing myeloid cells inhibit tumor growth *in vivo*.

## Materials and methods

### Cell lines and Mice

THP-1 cell line (Cat# TIB-202) was purchased from ATCC (American Type Culture Collection). Following human or mouse tumor cell lines were purchased from ATCC: NCI-H358 (Cat# CRL-5807), Hs578T (Cat# HTB-126), SK-MEL-28 (Cat# HTB-72), RM-1 (CRL-3310), 4T1(CRL-2539). In addition, MDA-MB-231 cell line was obtained from GeneCopoeia (Cat# SL018). THP-1 cells were cultured in RPMI-1640 medium containing 10% (vol/vol) FBS, 100 U/ml penicillin, 100 μg/ml streptomycin, and 50μM 2-mercaptoethanol (Thermo Scientific, Cat# 21985023). The tumor cells were cultured in complete medium (RPMI 1640 or DMEM) containing 10% (vol/vol) FBS, 100 U/ml penicillin, 100 ug/ml streptomycin.

Female Balb/c mice (stock no. 006584) between 6-7 weeks old were purchased from The Jackson Laboratory to serve as the syngeneic recipients for 4T1 tumor cell implantation and adoptive myeloid cell therapy. The mice, 5 per cage, were housed in the laboratory of Washington Biotechnology Inc. (Baltimore, MD) in autoclaved solid floor polycarbonate cages with filter-top, supplied with autoclaved bedding, at 22°C with a 12 hours’ light/dark cycle. Balb/c mice of the same sex and age were used to serve as the donor for bone marrow hematopoietic stem cells (HSCs) to generate immature myeloid cells for adoptive cell therapy. All animal experiments were approved by the Institutional Animal Care and Use Committee (IACUC) and conducted in accordance with the Washington Biotechnology, Inc. guidelines.

### Lentiviral vectors and lentiviral transduction

VSV-G pseudotyped third-generation lentiviral vectors encoding for CARIRs were custom ordered from VectorBuilder. Lentiviral vectors encoding human (Cat# LTV0746) or mouse PD-L1 (Cat# LTV1858) containing stable selection marker were purchased from G&P Biosciences. Lentiviral transduction of CARIR or PD-L1 were done in 24-well plate at MOI of 3, or as indicated, in the presence of 1× LentiBOOST (SIRION Biotech SB-P-LV-101-11). Transduction was facilitated through spinoculation by centrifuging the plate at 2500 RPM for 2h at 32°C. For overexpressing PD-L1, 24 hours following transduction, the cells were cultured in the presence of 2 μg/mL of puromycin for 10 days before analyzing PD-L1 expression via flowcytometry.

### Generation of primary myeloid cells or macrophages from human and mouse tissues

For generating human primary macrophages, human mobilized peripheral blood derived CD34^+^ cells (Lonza, 4Y-101C) were expanded in StemSpan SFEM II (StemCell Technologies, 09655), supplemented with 100 ng/mL human recombinant SCF, FLT3-L, TPO (Peprotech, HHCS3), 100 U/ml penicillin, and 100 ug/ml streptomycin. Macrophages were differentiated from the expanded CD34^+^ cells by first culturing for 7 days in StemSpan SFEM II supplemented with StemSpan Myeloid Expansion Supplement II (StemCell Tech 02694), and then culturing for another 7 days in IMDM media containing 20 ng/mL MCSF (PeproTech, 300-25) and 10% human AB serum (Sigma, H4522).

Mouse myeloid cells were prepared from bone marrow lineage^-^ cells of Balb/C mice. After isolated from the dissected tibia, fibula, and femur bones, the bone marrow cells were stained with biotin mouse lineage panel (BD Biosciences, 559971), followed by anti-biotin microbeads (Miltenyi, 130-105-637), before negative selection of lineage^-^ cells using AutoMACS (Miltenyi). Lineage^-^ cells were expanded in IMDM (Cytiva, SH30228.FS) supplemented with 20% BIT9500 (StemCell Tech, 09500), 100 ng/mL of murine recombinant SCF, FLT3-L, TPO (Peprotech, MHCS3), 100 U/ml penicillin, and 100 ug/ml streptomycin. Mouse myeloid cells were differentiated in IMDM (Cytiva, SH30228.FS) supplemented with 20% BIT9500 (StemCell Tech, 09500), 20 ng/mL M-CSF (Peprotech, 315-02), 100 U/ml penicillin, and 100 μg/ml streptomycin.

### Flow cytometry

The following monoclonal antibodies (mAbs) and their isotype controls were used for flow cytometry: APC human PD-1, APC human PD-L1, Brilliant Violet human PD-L1, APC human CD86, APC human CD11b, PE human CD247 (CD3z), APC mouse CD11b; PE human EGFR from R&D Systems; APC mouse PD-L1 from Tonbo Biosciences. In addition, following reagents were used in flow cytometry: recombinant human PD-L1 Fc chimera Biotin protein (R&D Systems); APC Streptavidin (Tonbo Biosciences); Zombie NIR viability dye from BioLegend; CellTrace Violet, CellTrace CFSE, and CellTrace Yellow from Invitrogen.

For flow cytometry analysis, cells were resuspended in Cell Staining Buffer (BioLegend) containing Fc receptor blocker (Miltenyi Biotec) and incubated for 10 minutes at 4°C. Then the cells were stained in PBS containing 1:1000 diluted Zombie NIR viability dye (BioLegend) for 30 minutes at room temperature. Finally, the cells were stained with antibodies diluted in Cell Staining Buffer for 20 minutes at 4°C. Cells were washed twice with Cell Staining Buffer between the staining steps and once prior to data acquisition using SONY SA3800 or SONY iD7000 spectral flow cytometer. The flow cytometry data was analyzed using FlowJo software.

### ELISA

Supernatant were collected at 72h following the co-culture of the engineered THP-1 macrophages and RM-1 target cells. The levels of human TNF-α, IL-1 β, and IL-6 were measured by ELISA using the DuoSet ELISA Kits (DY210-05, DY201-05, and DY206-05) from R&D Systems per the manufacturer’s protocol. The quantification was based on the OD values at 450 nm measured by a microplate reader (Multiskan Skyhigh, Thermo Scientific), subtracted by the readings at 540 nm.

### *In vitro* phagocytosis assay

Prior to initiating the coculture, macrophages were labeled with CellTrace Violet (Invitrogen C34557) and target cells were labeled with 1:1000 diluted CellTrace CFSE (Invitrogen C34554) or CellTrace Yellow (Invitrogen C34567) for 30 minutes at 37 °C. In some conditions, macrophages were pre-treated with 2μM cytochalasin D (Cayman Chemical 11330), 10μg/mL anti-PD1 antibody (BioXCell SIM 0010) or 10μg/mL human IgG4 isotype control antibody (BioXCell CP147) prior to co-culture with target cells. Macrophages and target cells were cocultured at a 5:1 E:T ratio for 3 hours at 37 °C in ultra-low attachment 96-well plates (Corning 7007), unless otherwise indicated. Cells were stained for viability with Zombie NIR (BioLegend 423106) and in some experiments with APC-PD-1 antibody (BioLegend 329908). Data was acquired on SONY SA3800 spectral flow cytometer and analyzed using FlowJo software.

### Flow cytometry-based cytotoxicity assay

To evaluate CARIR-mediated killing activity on PD-L1^+^ target cells, non-modified or CARIR engineered THP-1 cells (2.5 × 10^4^) were co-cultured with either RM-1^hPD-L1^ or RM-1 target cells (5 × 10^3^) in the presence of 5ng/ml PMA in 96-well round-bottom culture plate for 3 days. Following the co-culture, the cells were detached by Accutase (Stemcell Technologies) treatment at 37°C for 5 minutes. The cells were stained with Brillian Violet 605 anti-human PD-L1, APC anti-human CD11b, and Zombie NIR viability dye. Absolute counting beads (BioLegend) were added before data acquiring using SONY iD7000 spectral flow cytometry. The data was analyzed using flowjo software, and the percentage and absolute number of the remaining live tumor cells following the co-culture were compared between the groups.

### Subcutaneous 4T1 Tumor mouse model

Therapeutic effect of the engineered myeloid cells was tested in syngeneic Balb/c mice bearing subcutaneously implanted 4T1 breast cancer. The experiment was conducted in a blinded manner. On day -10, female Balb/c mice were subcutaneously injected on right flank with 5×10^4^ 4T1 T cells in 0.1ml PBS containing 20% Matrigel (Cat# 356231, Corning). On day 0, after the tumor became palpable, 24 of the mice were randomized into 3 groups (n = 8) based on tumor size. Each cage houses one mouse from each of the groups. On day 0, 7, and 14, the mice were treated intravenously through tail vein with 0.2ml PBS containing 1×10^7^ unmodified (WT-M) or CARIR engineered (CARIR-M) mouse bone marrow-derived myeloid cells, or PBS vehicle control. Body weight and tumor growth of the mice were measured 2-3 times a week. A digital caliper (Cat# 500-196-30, MSI Viking) was used for measuring the tumor size, and the tumor volume was calculated using the formula: Tumor volume = 1/2 (width × width × length). The study endpoints include tumor size beyond 2,000mm3, or when tumor growth causes more than 20% body weight loss, or when tumor becomes ulcerated.

### Statistics

Data for *in vitro* studies are representative of a minimum of 2 independent repeats, unless otherwise noted. The *in vitro* data are shown as mean ± SEM, with technical replicates plotted as individual data points. Data for *in vivo* study are shown as mean ± SEM with 8 mice per group. Statistical significance was determined by unpaired *t*-test or one-way ANOVA as indicated using GraphPad Prism software. For all statistical analysis, * indicating *p* < 0.05, ** *p* < 0.01, *** *p* < 0.001, and **** *p* < 0.0001.

## Results

### Stimulation through CARIR causes upregulation of co-stimulatory molecules and production of proinflammatory cytokines in human monocytic THP-1 cells

To direct macrophages to target the immune checkpoint receptor PD-L1, which is commonly expressed in many types of solid cancers^9^, we designed a CAR-like immune receptor (CARIR), where the extracellular domain is derived from the PD-1 ligand instead of a single chain fragment variable (ScFv) antibody. The intracellular signaling domain is derived from CD3ξ (CARIR-z) like a typical 1^st^ generation CAR design. A CARIR vector without the CD3ξ signaling domain was generated to serve as a control (CARIR-Δz) (**Figure 1A**). Lentiviral vectors were constructed to efficiently deliver the CARIR transgenes into human monocytic THP-1 cells. As shown in **Figure 1B**, 76.5 and 96.5% transduction efficiency were achieved at MOI of 3 for CARIR-Δz and CARIR-z, respectively, determined by staining for PD-1. A truncated version of EGFR (tEGFR) was incorporated in the lentiviral vector to serve as a potential safety module and kill switch, which could be detected in either CARIR-Δz and CARIR-z transduced THP-1 cells (**Suppl. Fig. 1A**). In addition, CD3z expression is confirmed in the CARIR-z transduced THP-1 cells (**Suppl. Fig. 1B**). Next, we confirmed that surface expressed CARIR in CARIR-z engineered THP-1 cells binds human PD-L1 following incubation with biotinylated recombinant human PD-L1 (**Suppl. Fig. 2**).

**Figure 1.**
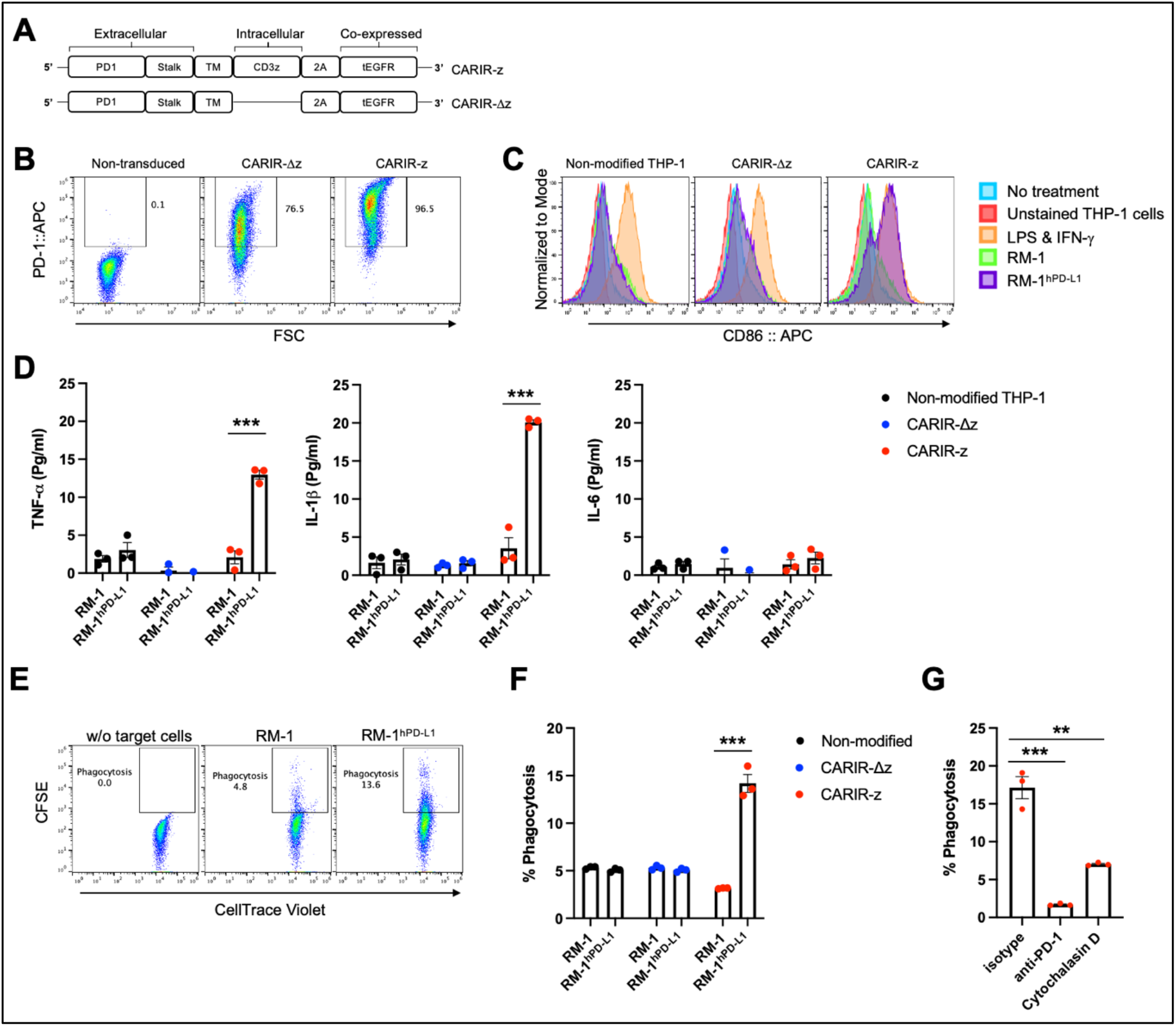
Functional expression of CARIR in human monocytic THP-1 cells. ***A***. Schematic illustration of lentiviral vector-encoded CARIR transgenes, with (CARIR-z) or without an intracellular signaling domain derived from human CD3ζ (CARIR-Δz). ***B***. CARIR expression in transduced human monocytic THP-1 cells based on flow staining of human PD-1. THP-1 cells were transduced with lentiviral vector encoding PD-1 CARIR at MOI of 3, and flow analysis was conducted 2 days following the lentiviral transduction. ***C***. The histogram shows the upregulation of co-stimulatory molecule CD86 in CARIR-z transduced THP-1 macrophages following co-culture with RM-1^hPD-L1^ cells. Non-transduced (non-modified) or CARIR transduced THP-1 cells were stimulated with 1ng/ml PMA for 24 hours, followed co-culture for 3 days as indicated. The CD86 expression was analyzed by flow cytometry, and the cells were gated on THP-1 cells based on the characteristic FSC/SSC parameters of the cells. ***D***. Cytokines levels measured by ELISA in the culture supernatant of the experiment as described in Fig. 1C above. ***E***. Representative flow cytometry dot plots depicting the phagocytosis events appeared in the Q2 quadrant, which is double positive for CellTrace CFSE labeled target cells and CellTrace Violet labeled THP-1 macrophages. Non-modified or CARIR transduced THP-1 cells were treated with 5ng/ml PMA for 24 hours to differentiate the cells toward macrophages, followed by co-culture with RM-1 or RM-1^hPD-L1^ at E:T ratio of 5:1 for 3 hours. Both the effectors and the target cells were fluorescent dye labeled before setting up the co-culture. The cells were gated on FSC/SSC, singlets, and live THP-1 macrophages (CellTrace Violet^+^). ***F***. Bar graph summarizes the % phagocytosis during the 3-hour co-culture, in experiment as described for Fig. 1F. ***G***. BAR graph shows the inhibition on phagocytosis in the presence of anti-PD-1 blockade antibody or cytochalasin D in experiment as described for Fig. 1F. The data were expressed as mean ± SEM. **p < 0.01 and \*\*\**p* < 0.001, by non-paired student t test with 2-tailed distribution.

**Figure 2.**
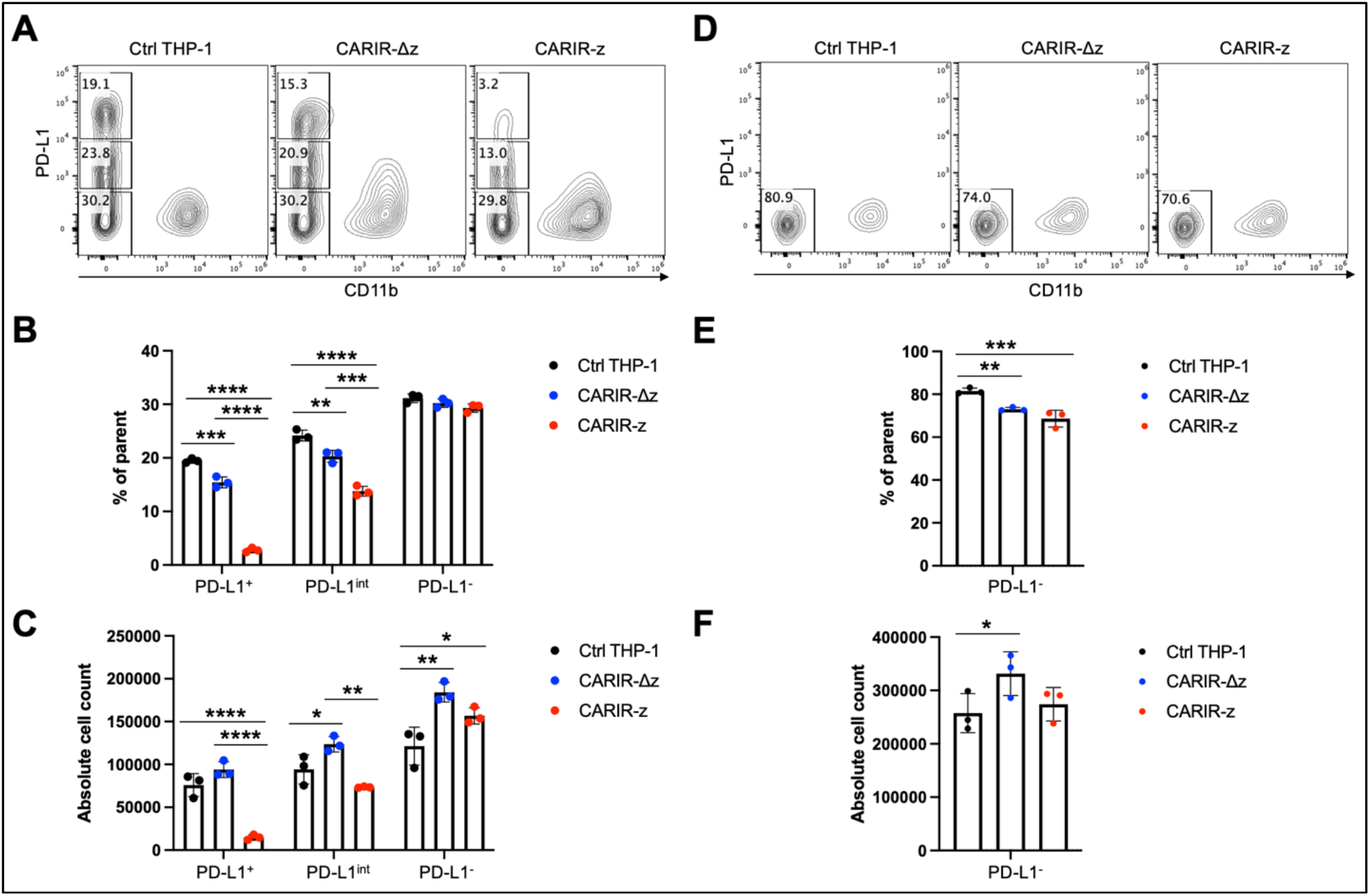
CARIR expression in THP-1 macrophages enabled significant cytotoxicity against PD-L1^+^ target cells. Non-modified-(Ctrl), CARIR-Δz, or CARIR-z-engineered THP-1 effector cells were co-cultured in the presence of 5ng/ml PMA for 3 days with RM-1^hPD-L1^ or WT-RM-1 target cells at effector to target ratio of 5:1. Following the co-culture, the cells were stained with APC anti-human CD11b, Brilliant Violet 605 anti-human PD-L1, and Zombie NIR viability dye. The number of the remaining live CD11b^-^ target cells following the co-culture was quantified by flow cytometry with the use of absolute counting beads. ***A***. Representative contour plots show the percentage change for PD-L1^+^, PD-L1^int^, and PD-L1^-^ RM-1^hPD-L1^ tumor cells following the co-culture. ***B***. BAR graphs summarize the percentage data shown on panel A. ***C***. BAR graphs summarize the absolute cell count data shown on panel A. ***D***. Representative contour plots show the percentage change of RM-1 tumor cells (PD-L1^-^) following the co-culture. ***E***. BAR graph summarizes the percentage data shown on panel D. ***F***. BAR graph summarizes the absolute cell count data shown on panel D. Tumor cells were gated on non-beads, live, singlets, and CD11b^-^. Data were presented as mean ± SEM. \**p* < 0.05, \*\**p* < 0.01, \*\*\**p* < 0.001, and \*\*\*\**p* < 0.0001 by one-way ANOVA.

**Figure 3.**
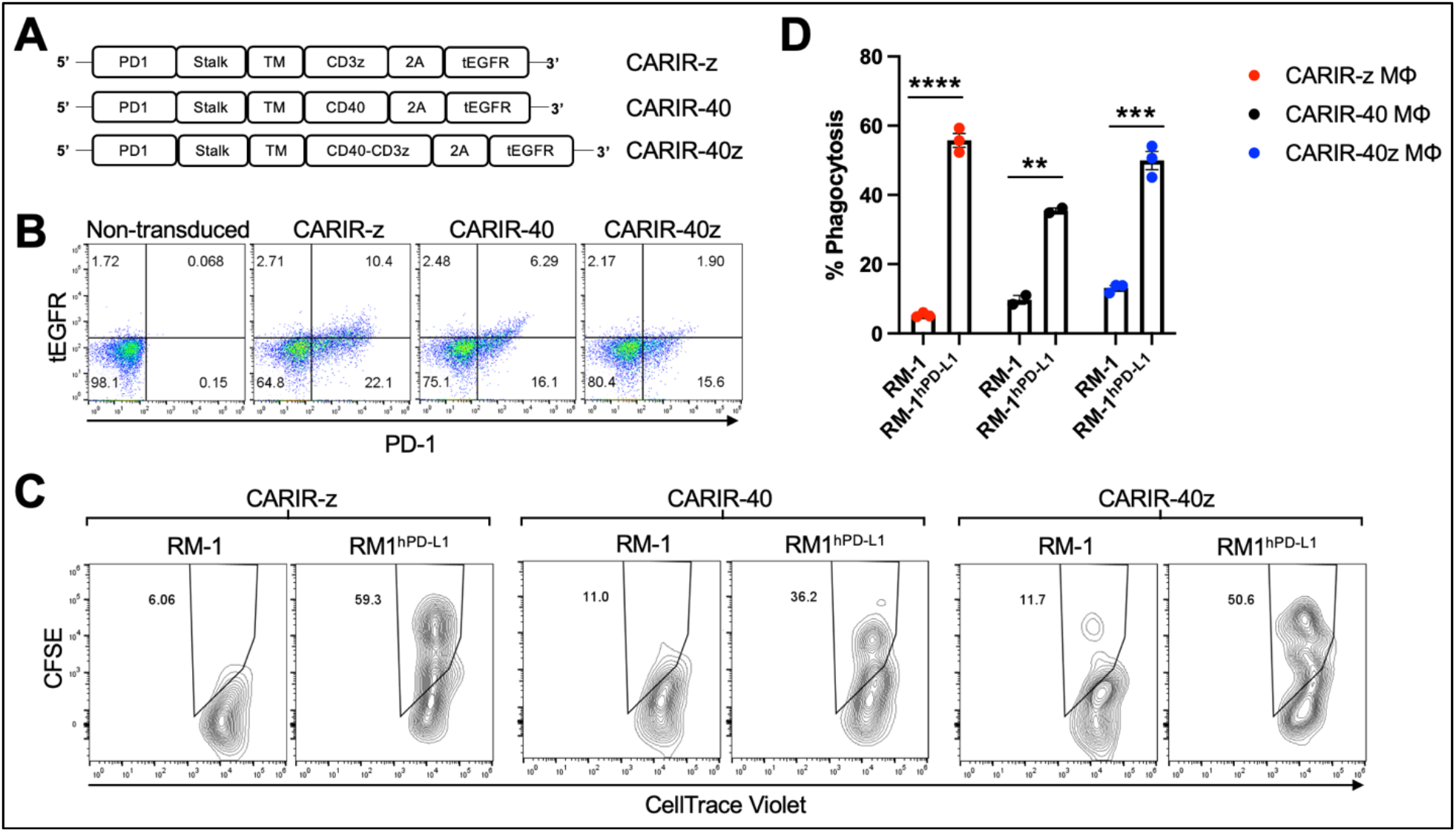
CARIR expression in primary human macrophages enhanced phagocytosis against human PD-L1^+^ target cells. ***A***. Schematic illustrations show different CARIR constructs varying in the intracellular signaling domain(s). ***B***. Flow dot plots show the efficiency of CARIR transduction in human HSCs based on staining for PD-1. ***C***. Representative flow plots showing the % phagocytosis of CARIR expressing primary human macrophage against RM1 or RM1^hPD-L1^ target cells. For generating CARIR modified human macrophages to serve as the effector cells, human CD34^+^ hematopoietic stem cells (HSCs) were engineered to express CARIR through lentiviral transduction, and then differentiated into macrophages. Fluorescent dye labeled effector cells and target cells were co-cultured for 3 hours before flow analysis for the % phagocytosis. The cells were gated on FSC/SSC, singlets, live, and PD-1^+^Violet^+^ macrophages. The CellTrace Violet and CFSE double positive population represent macrophages that have phagocytosed target cells. ***D***. The bar graph summarizes the flow data shown in panel C. \*\**p* < 0.01, \*\*\**p* < 0.001, and \*\*\*\**p* < 0.0001 by unpaired student *t* test with two tailed distributions.

The surface expressed CARIR is functional, as stimulation with plate-bound anti-PD-1 led to an upregulation of co-stimulatory molecules CD86 and CD80 in CARIR-z-THP-1 cells (**Suppl. Fig.3**). To facilitate the functionality test for CARIR modified macrophages, we generated a surrogate PD-L1-expressing model system by stably transducing murine RM-1 cells with human PD-L1 (RM-1^hPD-L1^) (**Suppl. Fig. 4**). Next, we differentiated unmodified, CARIR-Δz, and CARIR-z transduced THP-1 cells towards macrophages through PMA stimulation for 24 hours followed by co-culture with WT RM-1 or RM-1^hPD-L1^ cells. Following 3 days of co-culture, THP-1 cells were analyzed for expression of the co-stimulatory molecule CD86 by flow cytometry. As expected, following 3 days of co-culture, CD86 was upregulated in CARIR-z expressing THP-1 cells but not in CARIR-Δz expressing THP-1 cells. Co-culture with unmodified RM-1 cells did not lead to significant change of CD86 expression (**Fig. 1C**). In addition, we analyzed the prototypical M1 inflammatory cytokines TNF-α, IL-1Δ and IL-6 in the supernatant from the co-culture. The production of proinflammatory cytokines, including TNF-α and IL-1Δ, were significantly increased in CARIR-z expressing THP-1 (**Fig. 1D**).

Next, we sought to determine if activation through CARIR-z enhances the macrophage phagocytic activity. To do this, unmodified, CARIR-Δz, and CARIR-z transduced THP-1 derived macrophages were co-cultured with either RM-1 or RM-1^hPD-L1^ for 3 hours before evaluation by flow cytometry for phagocytosis events (**Fig. 1E**). As expected, co-culture with RM-1 cells did not increase phagocytosis while co-culture with RM-1^hPD-L1^ led to a 2.8-fold increase (14.2 versus 5.1%) of the % phagocytosis against the target cells by the CARIR-z-THP-1 macrophages. In the contrast, CARIR-Δz expression in THP-1 macrophages did not increase phagocytosis against RM-1^hPD-L1^ target cells (**Fig. 1F**). As expected, there was no increase of % phagocytosis by CARIR-z expression in THP-1 macrophages in the presence of the actin polymerization inhibitor, cytochalasin D. Notably, the increase of phagocytosis against PD-L1^+^ target cells by CARIR-z-THP-1 macrophages were CARIR-dependent, since the increase in phagocytosis was completely obligated in the presence of monoclonal anti-PD-1 blockade antibody (**Fig. 1G**). Taken together, these results demonstrate that CARIR-mediated activation enhances phagocytosis and polarize macrophages towards a proinflammatory phenotype.

### CARIR expression enables specific killing of PD-L1^+^ target tumor cells by THP-1 macrophages

To determine if CARIR expression could direct macrophages to kill PD-L1^+^ tumor cells, A flow-based cytotoxicity assay was used. THP-1 macrophages were served as the effector cells and RM-1^hPD-L1^ or RM-1 tumor cells as the target cells. Since monocytic THP-1 cells are commonly differentiated into macrophages with PMA treatment, to streamline the assay, we co-cultured the effector cells and the target cells in the presence of PMA. PMA treatment had no significant effect on the viability of either RM-1^hPD-L1^ or RM-1 tumor cells (**Suppl. Fig. 5A**). In addition, the target tumor cells could be distinguished with the effector THP-1 cells due to a lack of expression of the myeloid marker CD11b, and RM-1^hPD-L1^ cells could be further divided into three populations: CD11b^-^PD-L1^+^, CD11b^-^PD-L1^int^ and CD11b^-^PD-L1^-^ (**Suppl. Fig. 5B**). Compared to control THP-1, co-culture with CARIR-z-THP-1 led to a significant loss of the PD-L1^+^, but not PD-L1^int^ or PD-L1^-^ RM-1-^hPD-L1^ cells. In the contrary, CARIR-Δz-THP-1 significantly increased the PD-L1^int^ and PD-L1^-^ RM-1-^hPD-L1^ cells proliferation (**Fig. 2A - C**). The lack of cytotoxicity of CARIR-z-THP-1 on PD-L1^-^ target cells was confirmed when non-modified RM-1 cells were served as the target cells (**Fig. 2D - E**). These results indicate that the CD3 zeta signaling domain is necessary, and there is a threshold on PD-L1 expression levels for CARIR-mediated killing by the engineered THP-1 macrophages.

**Figure 4.**
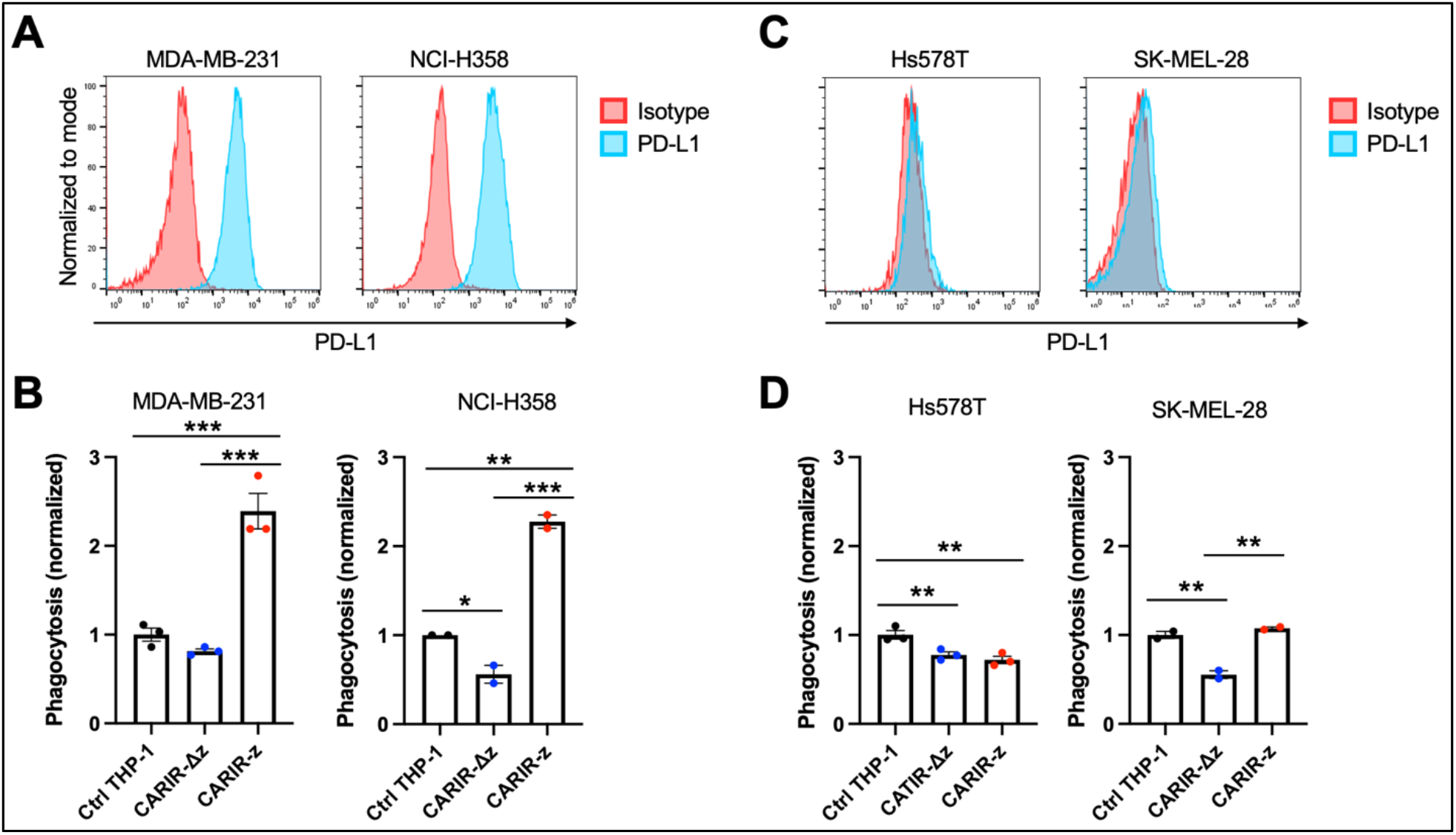
CARIR expression in human THP-1 macrophages increased phagocytosis against PD-L1^+^ human tumor cells. ***A & C***. Histogram shows the detection of cell surface PD-L1 expression by flow cytometry in cultured human tumor cell lines. ***B & D***. Phagocytosis activity against PD-L1^+^ tumor cell lines MDA-MB-231 and NCI-H358 (B), or PD-L1^-^ tumor cell lines Hs578T and SK-MEL-28 (D). Human monocytic THP-1 cells were either nonmodified (Ctrl THP-1) or engineered to express either CARIR-Δz or CARIR-z through lentiviral transduction, and then differentiated toward macrophages by culturing in the presence of 1 ng/ml PMA for 24 hours. CellTrace Violet labeled THP-1 effectors were co-cultured with CellTrace Yellow labeled indicated tumor cells for 4 hours, followed by flow cytometry analysis of the % phagocytosis. Cells were gated on live, singlets, and violet^+^ cells. The events that were double positive for CellTrace Violet and CellTrace Yellow were considered as phagocytic events. The phagocytosis activity was normalized to the non-modified THP-1 condition. The data was presented as mean ± SEM. \**p* < 0.05, \*\**p* < 0.01, and \*\*\**p* < 0.00 by one-way ANOVA.

**Figure 5.**
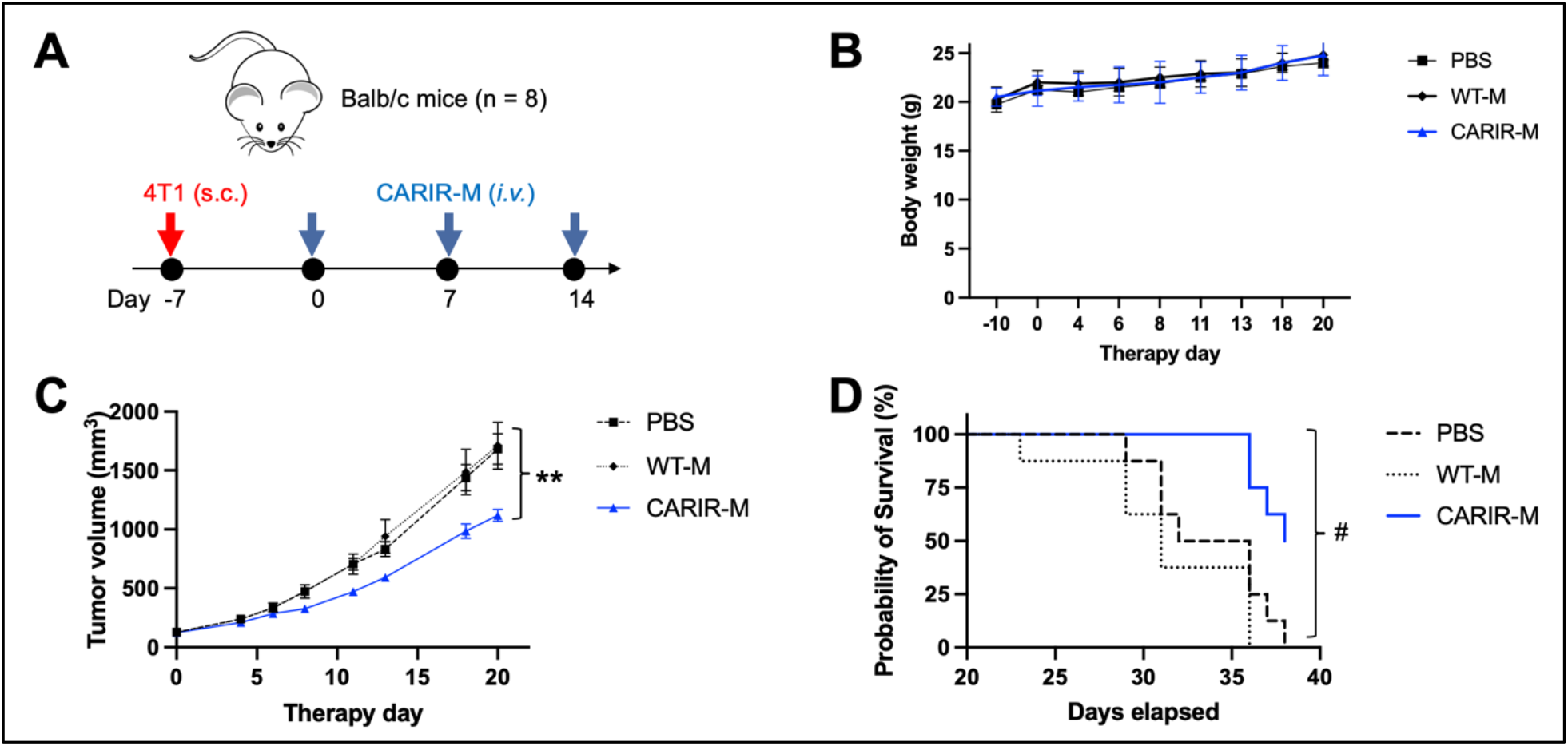
Adoptive transfer of CARIR modified myeloid cells (CARIR-M) slows 4T1 tumor growth and prolongs survival. ***A***. Schematic timeline for the animal experiment. Syngeneic Balb/c mice were subcutaneously implanted with 5 × 10^4^ 4T1 breast cancer cells on day -7. Starting on day 0, the mice were injected through the tail vein with 3 weekly doses of 10 × 10^6^ either CARIR-M or control non-modified myeloid cells (WT-M). An additional control group of mice were treated with PBS. Body weight (**B**), tumor volume (**C**), and probability of survival (**D**) were measured 2-3 times per week. Data are presented as mean ± SEM. **p < 0.01: CARIR-M vs WT-IMC or PBS by type II ANOVA. #p < 0.05 (CARIR-M vs PBS) and p < 0.01(CARIR-M vs WT-M) by Gehan-Breslow-Wilcoxon test.

### CARIR expression in primary human macrophages increases phagocytosis against PD-L1^+^ target cells

Next, we tested the functionality of CARIR in primary human macrophages, prepared from mobilized peripheral blood derived CD34^+^ hematopoietic stem cells (HSCs). For CARIR, in addition to CARIR-z, we created additional lentiviral constructs with the intracellular signaling domain derived from CD40 (CARIR-40) or from both CD40 and CD3ζ (CARIR-40z) (**Fig. 3A**). HSCs were transduced with lentiviral vector for either CARIR-z, CARIR-40, or CARIR-40z. The transduction efficiencies were between 17 – 32% as determined by PD-1 expression 2 days following the transduction (**Fig. 3B**). The transduced cells were expanded and sequentially differentiated towards macrophages. The phagocytosis assay was performed by co-culturing these modified human primary macrophages with RM-1 or RM-1^hPD-L1^ target cells for 3 hours. Consistent to the result obtained with CARIR-z THP-1 macrophages, following 3 hours of co-culturing, CARIR-z modified human primary macrophages had a 10-fold increased phagocytosis against RM-1^hPD-L1^ than RM-1 targets (55.8% vs 5.2%). Similarly, CARIR-40 or CARIR-40z expression in human primary macrophages also led to significant increase in phagocytosis against RM-1^hPD-L1^ than RM-1 targets (35.5 vs 9.7% for CARIR-40; 49.9 vs 13.1% for CARIR-40z) (**Fig. 3C&D**). These results further confirmed the functionality of CARIR expression in macrophages in enhancing phagocytosis against human PD-L1^+^ target cells. In addition, the results indicated that CD3ζ signaling domain is sufficient for the CARIR-mediated functionality. Therefore, in the rest of the study we will be focusing on testing CARIR functionality in the format of CARIR-z configuration.

### CARIR expression in human THP-1 macrophages enhances phagocytosis against PD-L1^+^ human tumor cell lines

Next, instead of artificial human PD-L1 engineered target cells, we asked if PD-L1^+^ human solid tumor cells could be targeted by CARIR modified macrophages *in vitro*. To this goal, we utilized 4 human solid tumor cell lines, including MDA-MB-231, NCI-H358, Hs578T, and SK-MEL-28, to serve as the target cells. The former two lines are PD-L1^+^, while the other two lines were barely detectible for PD-L1 surface expression by flow cytometry (**Fig. 4A&C**). *In vitro* phagocytosis assays were performed using each of the above tumor cells as the targets and non-modified or engineered THP-1 macrophages as the effectors. CARIR-z THP-1 had significantly increased phagocytosis activity against both the PD-L1^+^ tumor cell lines, the triple negative breast cancer (TNBC) line MDA-MB-231 and non-small-cell lung cancer (NSCLC) line NCI-H358, as compared to either the nonmodified or CARIR-Δz-THP-1 macrophages conditions. In the contrast, CARIR expression in THP-1 macrophages did not lead to an increased phagocytosis activity against the PD-L1 negative Hs578T and SK-MEL-28 tumor cell lines (**Fig. 4B&D**).

### Adoptive transfer of CARIR modified myeloid cells ameliorated tumor growth in a syngeneic TNBC tumor mouse model

Since PD-1 CARIR is specific to PD-L1, CARIR modified macrophage approach can be considered as a cell therapy version of immune checkpoint therapy, enhanced by the chimeric receptor technology. Next, we sought to evaluate the anti-tumor functionality of CARIR *in vivo* using syngeneic Balb/c mice bearing aggressive 4T1 triple negative breast cancer (TNBC). 4T1 tumor expresses positive albeit low cell surface PD-L1 and normally does not respond to classic anti-PD-L1 checkpoint blockade^10^. Thus, we prepared a lentiviral vector encoding a murine analogue of human PD-1 CARIR-z. CARIR engineered myeloid cells were prepared by transducing donor Lin^-^ HSCs with the mouse version of CARIR-z. A 48% mouse PD-1 expression (CARIR transduction efficiency) was achieved at the MOI of 10. After a total of 6 days expansion in the presence of SCF, TPO and Flt3L, the transduced cells were further myeloid differentiated in M-CSF medium for 24 hours. In mice with established 4T1 tumor, weekly infusion of CARIR-z myeloid cells (CARIR-M) significantly slowed tumor growth and prolonged survival (**Fig. 5**). Of note, no significant differences were observed in body weight measurements between the CARIR-M, WT-M, or the non-treated (PBS) group, suggesting a lack of overt toxicity associated with CARIR-M treatment (**Fig. 5B**).

## Discussion

Macrophages are highly plastic in their functionality and have capacity not only in infiltrating solid tumor, but also in tumor cell phagocytosis and orchestrating adaptive immune response through tumor antigen presentation. Adoptive cell therapies using chimeric receptor engineered macrophages or its immediate precursor, monocytes, are becoming an exciting avenue in developing effective treatments for solid cancers^3,11,12^. However, devising a traditional CAR approach to target tumor-specific antigens remains a significant challenge as unique and broadly expressed tumor-specific antigens are rare. Moreover, solid cancers are heterogenous in nature and targeting a single tumor antigen often leads to antigen escape. As a common mechanism of immune evasion, solid tumor cells and/or tumor-infiltrating immune cells upregulate the immune checkpoint molecule PD-L1^13^. As a result, several studies have described targeting PD-L1 using CAR-T^14–18^ or CAR-NK^19^ as cell therapy strategies for cancer. To our knowledge, we have demonstrated in this study for the first time that myeloid cells or macrophages that modified to express CARIR could be used to effectively target PD-L1 in solid cancer.

As a chimeric receptor, CARIR is akin to the previously described PD-1-CD28 fusion receptor or PD-L1-specific chimeric switch receptor (CSR) for T cells ^20–22^. The common feature of these chimeric receptors is that the extracellular domain is derived from the immune checkpoint molecule PD-1. However, the intracellular domain usage and the functional utility of these chimeric receptors differ. The only cytosolic domain of PD-1-CD28 fusion receptor or PD-L1-specific CSR is derived from the costimulatory molecule CD28, and the purpose of expressing these chimeric receptors is to enhance the functionality of tumor-specific TCR-T or CAR-T cells^20–22^. In the present study, however, the purpose of the CARIR is to direct macrophages to recognize and attack PD-L1^+^ solid cancer cells and reprogram immunosuppressive TME. Thus, the intracellular region of CARIR is derived from the cytosolic signaling domain(s) of CD3z and/or CD40. As expected, CARIR-z, but not CARIR-Δz, induced a proinflammatory phenotype of THP-1 macrophages and increased phagocytosis and killing activities against RM-1^hPD-L1^ target cells, whereas no effect was seen against WT RM-1 (**Fig. 1&2**). Although exclusively expressed in T cell lineages, the signaling domain of CD3z in a chimeric receptor format is known to be able to transmit signal and trigger phagocytosis in macrophages^11^. In this study we also tested a CARIR version with the macrophage specific CD40 signaling domain. However, based on phagocytosis efficiency of PD-L1^+^ target cells (**Fig 3 C&D**), CD3z outperformed CD40 and CD40/CD3z, which let us to focus our study on the CARIR-z version. It is of note, high expression of PD-L1 has also been found in myeloid derived suppressor cells (MDSC) and subsets of TAMs^7^, making them all attractive targets in treating solid cancer^24^.

Under physiological condition, PD-1/PD-L1 signaling functions as a mechanism for maintaining immune tolerance, preventing excess immune cell activity that can lead to autoimmunity and tissue damage. This immune-inhibiting axis is exploited by many types of malignancies to evade the body’s anti-tumor immune response. Targeting PD-L1 tilts the immune homeostasis from immune tolerance towards anti-tumor cytotoxicity. As seen with the remarkably successful immune checkpoint therapies, certain level of off-tumor cytotoxicity is inevitable and is expected for CARIR modified macrophages therapy as well. However, there are a few reasons that may argue against a major safety concern for the application of CARIR, which utilizes natural PD-1 domain instead of ScFv antibody for target recognition: 1) the affinity between human PD-1 (CARIR) and its ligand PD-L1 is about 7.2 μM^25^, which is several logs lower than clinical approved anti-PD-1/PD-L1 monoclonal antibodies^26^, suggesting CARIR modified macrophages might be able to better discriminate between upregulated PD-L1 in the case of solid cancer and the physiological expression of PD-L1 in normal healthy cells; 2) the result from our flow-based killing assay in Figure 2 indicates a threshold on PD-L1 expression levels on target cells may exist for CARIR-mediated killing activity by the engineered THP-1 macrophages; and 3) when tested in *in vivo* in fully immunocompetent syngeneic 4T1 tumor mouse model (**Fig. 5**), the adoptive transfer of a murine analog of CARIR modified myeloid cells significantly slowed the aggressive 4T1 tumor growth and prolonged survival, but no change of body weight was observed during the experiment as compared to the control groups, suggesting a lack of overt treatment toxicity.

Taken together, we described here an approach employing CARIR modified macrophages as a potential treatment for PD-L1^+^ solid cancer. CARIR expression directed macrophages towards PD-L1^+^ target cells, led to macrophages activation, increased phagocytosis and killing of PD-L1^+^ target cells, potentiate tumor antigen cross-presentation, and adoptive transfer of CARIR transduced myeloid cells significantly protected fully immunocompetent syngeneic mice of aggressive 4T1 tumor. These proof-of-principle results support the utility of CARIR modified autologous macrophages to be developed as a potential therapy for PD-L1^+^ solid cancer.

## Acknowledgements

The authors thank Dr. Amir Saberi, Dr. Jeff Harding and members of the Vita team for technical assistance and construct designs; Dr. Yinghua Wang (Washington Biotechnology, Inc.) for helpful discussions and technical assistance on mouse work. Preprint of this article has been deposited in BioRxiv.

## Contributions

KMC and DG designed, conducted experiments, interpreted data, and edited the manuscript. BG and MM performed experiments. SV designed, performed experiments, and interpreted data. AHZ and PA designed, interpreted data, supervised the work, and wrote the manuscript.

## Competing interests

Authors (KMC, DG, BG, SV, MM, AHZ, and PA) are employees and/or shareholders of Vita Therapeutics, Inc. AHZ, PA, KMC, and DG are inventors of pending patent(s) involving the generation and use of CARIR-modified myeloid cells for treating cancer.

## List of Abbreviation

CAR: (Chimeric Antigen Receptor)
CARIR: (CAR-like Immune Receptor)
PD-1: (Programmed Cell Death Protein 1)
PD-L1: (Programmed Cell Death Ligand 1)
TME: (Tumor Microenvironment)
tEGFR: (Truncated Epidermal Growth Factor Receptor)
TAMs: (Tumor Associated Macrophages)
HSCs: (Hematopoietic Stem Cells)
IACUC: (Institutional Animal Care & Use Committee)
mAb: (Monoclonal Antibody)
ELISA: (Enzyme-Linked Immunosorbent Assay)
SEM: (Standard Error)
ScFv: (Single Chain Variable Fragments)
MOI: (Multiplicity of Infection)
PMA: (Phorbol Myristate Acetate)
APC: (Antigen Presenting Cells)
TNBC: (Triple Negative Breast Cancer)
CSR: (Chimeric Switch Receptor)
MDSC: (Myeloid Derived Suppressor Cells)
Ctrl: (Control)

