## Supplemental Figures for "Targeting PD-L1 in solid cancer with myeloid cells expressing a CAR-like immune receptor"

**Supplemental Material**

**Supplemental Figure 1**

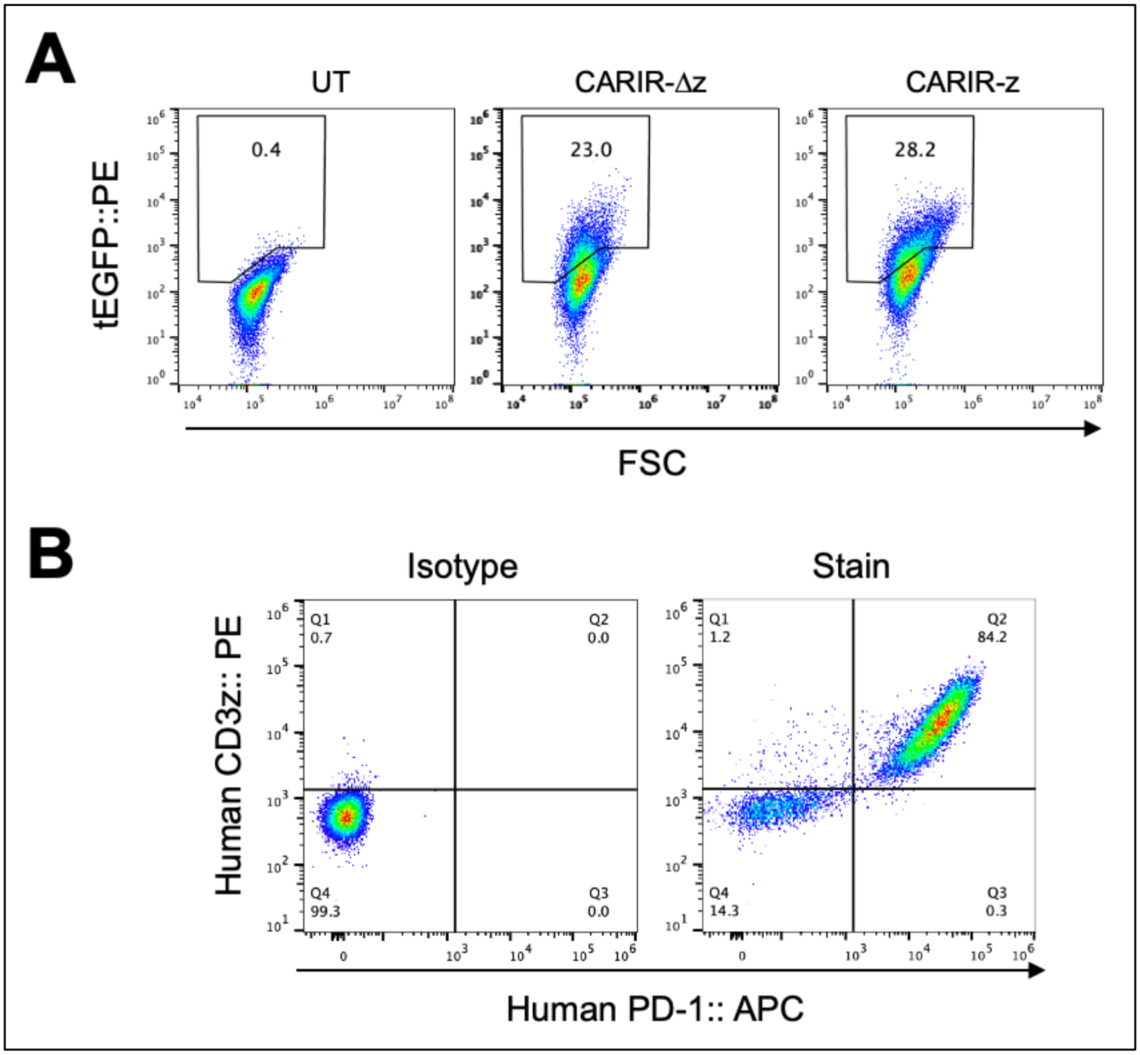


**Suppl. Fig. 1. Expression of a truncated EGFR (tEGFR)/kill switch and CD3z in CARIR transduced human monocytic THP-1 cells**. THP-1 cells were not-transduced (UT) or transduced with the lentiviral vector encoding CARIR-Δz or CARIR-z at a MOI of 3. The cells were stained with the indicated fluorescence labeled antibodies and analyzed by flow cytometry. ***A***. Pseudo-color plots show the surface detection of truncated EGFR in the CARIR-Δz and CARIR-z engineered THP-1 cells. ***B***. Pseudo color plots show the detection of CD3zeta (CD3z) through intracellular staining in the CARIR-z transduced THP-1 cells. The cells were gated on size and then live and singlets.

### Supplemental Figure 2

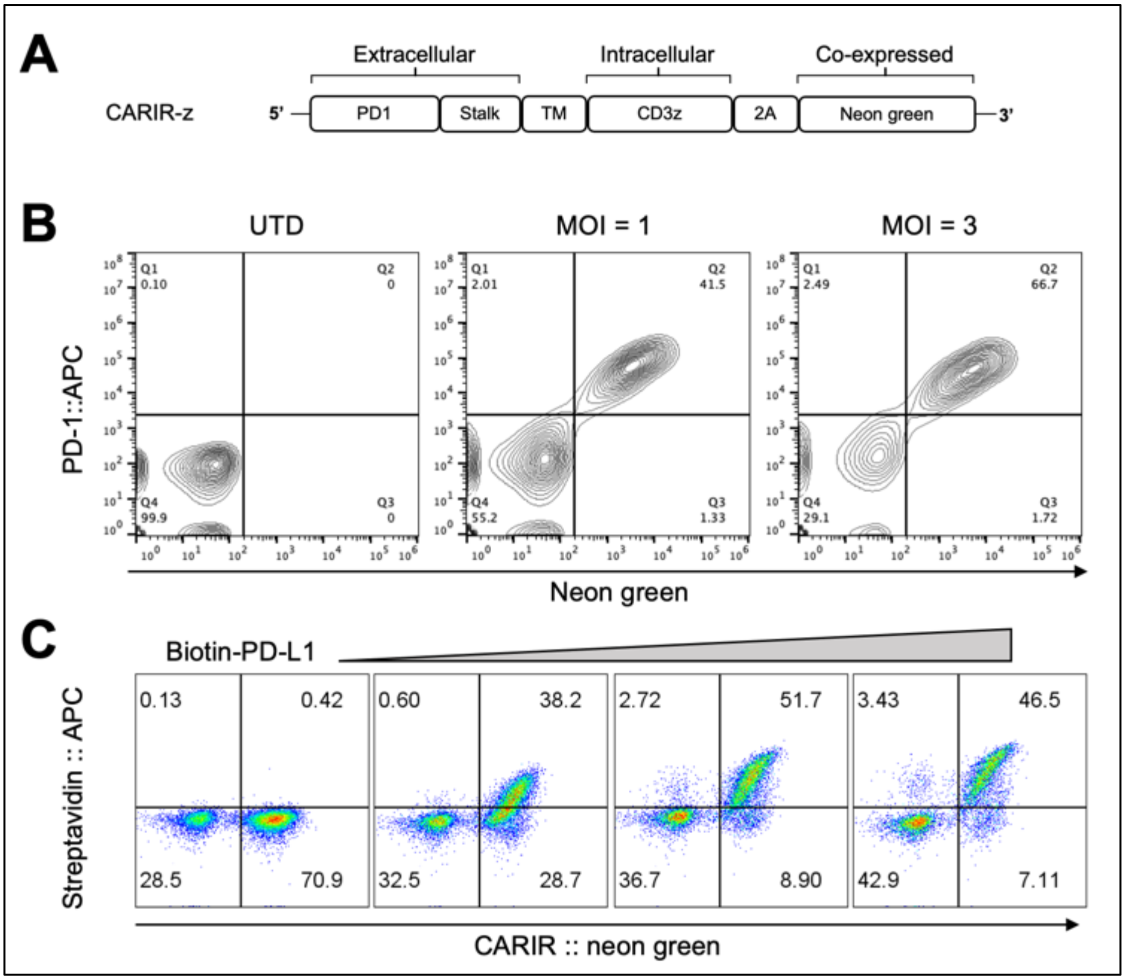


**Suppl. Fig. 2. CARIR expressed on transduced human monocytic THP-1 cells binds with human PD-L1 ligand**. ***A.*** Diagram for a CARIR-z lentiviral vector that encoding a co-expressed Neon green marker. ***B.*** Co-expression of PD-1 and Neon green marker in transduced human monocytic THP-1 cells, as measured by flow cytometry. ***C.*** PD-1 CARIR expressed on transduced THP-1 cells binds biotinylated human PD-L1. CARIR transduced THP-1 cells were stained without or with increasing amount of biotinylated human PD-L1, followed by staining with streptavidin APC.

### Supplemental Figure 3

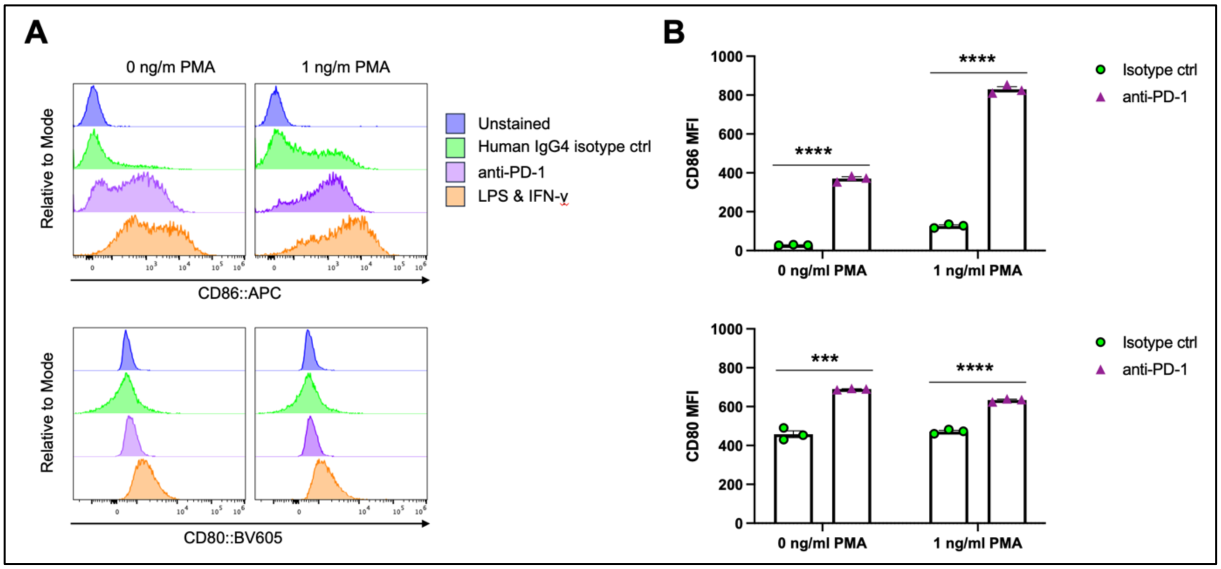


**Suppl Fig 3. CARIR-mediated upregulation of co-stimulatory molecules CD86 and CD80 in CARIR-z-THP-1 cells.** The CARIR-z-THP-1 cells were pretreated without or with 1 ng/ml PMA for 2 days in regular tissue culture flask. After resting for 8 hours in the absence of PMA, the cells ($1 \times 10^5$ per well) were added to a 96-well U-bottom culture plate that has been pre-coated with 2μg/ml anti-human PD-1 pembrolizumab biosimilar (anti-PD-1) or human IgG4 isotype control (isotype ctrl). Two days later, the cells were analyzed by flow cytometry for the expression of co-stimulatory molecules CD86 and CD80. ***A.*** The off-set histograms show the expression of CD86 and CD80. ***B.*** The bar graphs summarized the data between the anti-PD-1 and human IgG4 isotype control groups shown on panel A. MFI: median fluorescence intensity. The experiment was conducted in triplicate, and the data was presented as mean ± SEM. $^{**}p < 0.01$, $^{***}p < 0.001$, and $^{****}p < 0.0001$ between anti-PD-1 and human IgG4 isotype control groups by two-tailed student $t$ test.

**Supplemental Figure 4**

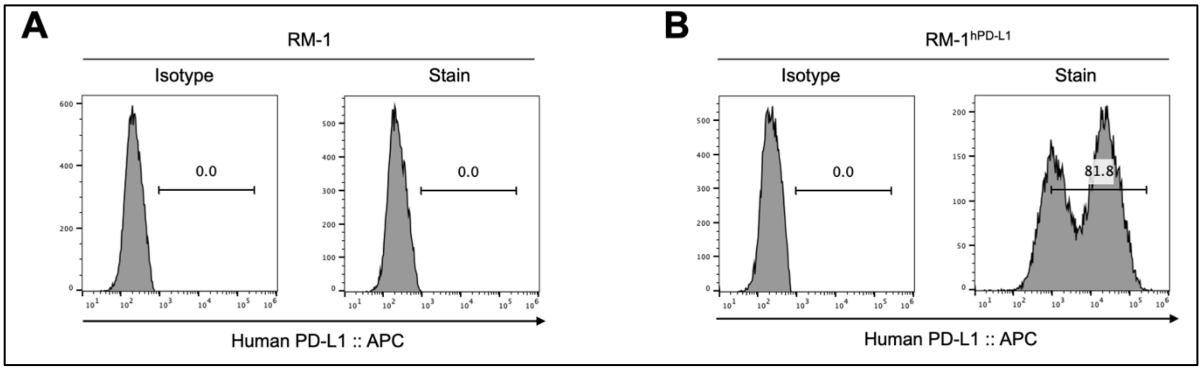


**Suppl. Fig. 4. Human PD-L1 expression in the generated RM-1hPD-L1, as compared to WT RM-1.** RM-1 cells were lentivirally transduced to overexpress human PD-L1. ***A.*** Histogram shows that RM-1 cells are stained negative for human PD-L1. ***B.*** Histogram shows that majority of the RM-1hPD-L1 cells are stained positive for human PD-L1 expression.

### Supplemental Figure 5

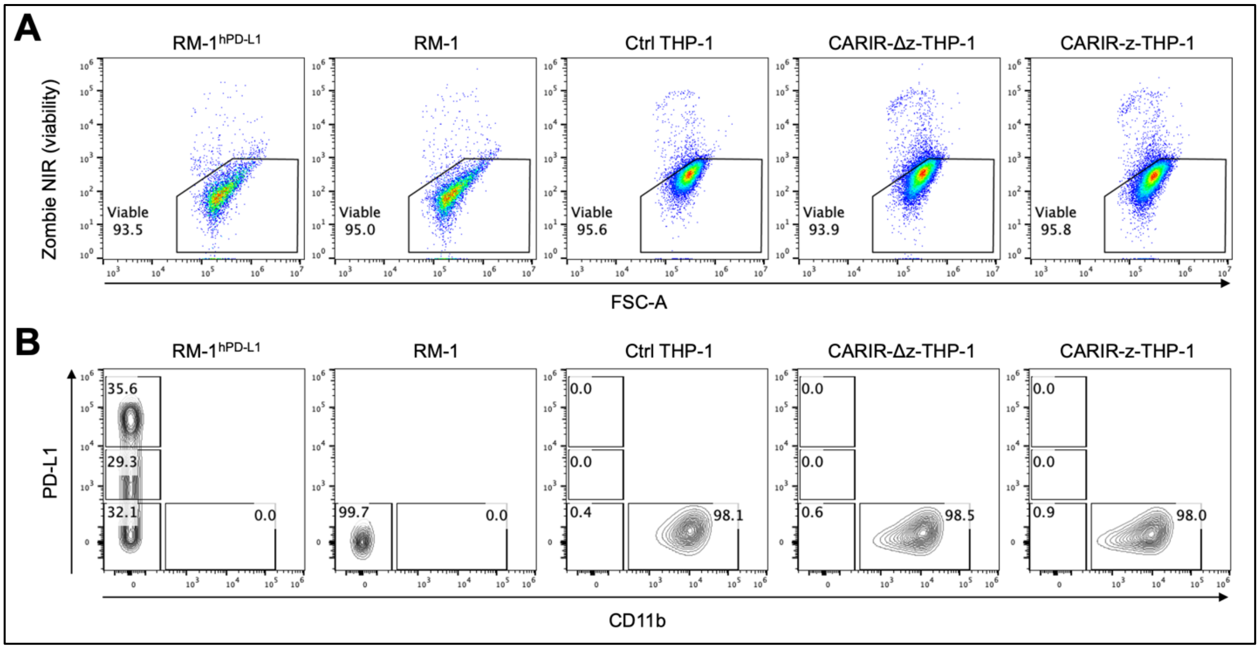


**Suppl. Fig. 5. Culturing in the presence of 5ng/ml PMA did not affect the viability of target RM-1$^{hPD-L1}$ or RM-1 tumor cells.** RM-1$^{hPD-L1}$, WT-RM-1, as well as effector cells, including Non-modified- (Ctrl), CARIR-Δz, or CARIR-z-engineered THP-1 cells, were cultured in the presence of 5ng/ml PMA for 3 days. Then the cells were stained with APC anti-human CD11b, Brilliant Violet 605 anti-human PD-L1, and Zombie NIR viability dye. ***A.*** Representative dot plots show viability of the cells. The percentage of viable cells are similar between the cell types. ***B.*** Representative contour plots show the differential expression of PD-L1 and CD11b among the target tumor cells or the effector THP-1 cells. Cells were gated on size and singlets.
